# Context-dependent functional diversity of dorsomedial posterior parietal neurons revealed by single-unit fMRI mapping during naturalistic viewing

**DOI:** 10.64898/2026.05.27.728098

**Authors:** Xufeng Zhou, Lei Wang, Shu K. E. Tam, Michael Ortiz-Rios, Dazhi Yin, Sze Chai Kwok

## Abstract

The dorsomedial posterior parietal cortex (dmPPC) plays an important role in episodic processing by integrating sensory, cognitive, and motor information across distributed brain systems. However, how individual dmPPC neurons participate in large-scale functional organization during naturalistic experience remains poorly understood. To address this question, we combined single-unit electrophysiology and awake fMRI in separate cohorts of rhesus macaques viewing identical naturalistic video stimuli. Using single-unit fMRI mapping, we generated whole-brain neuron–BOLD functional maps by correlating individual neuronal activity with voxel-wise fMRI signals across the brain. We found that neuron–BOLD functional maps exhibited strong context-dependent organization, with neurons recorded during the same video context showing substantially greater similarity than neurons recorded during different video conditions. Compared with neuronal spiking activity or critical fMRI frames alone, neuron–BOLD functional maps more robustly capture contextual structure. Despite this shared large-scale organization, neighboring neurons recorded simultaneously from the same electrode often displayed markedly distinct whole-brain association patterns, revealing substantial local functional heterogeneity within the dmPPC. In addition, distributed cortical and medial temporal regions exhibited highly context-dependent neuron–BOLD association patterns during naturalistic viewing. Together, these findings demonstrate that dmPPC neurons participate in dynamic and heterogeneous large-scale functional organization during naturalistic episodic processing. More broadly, this study establishes single-unit fMRI mapping as a framework for linking single-neuron activity to distributed whole-brain dynamics across contextual conditions.

**Significance Statement:** Using single-unit fMRI mapping, this study examined how individual dorsomedial posterior parietal cortex (dmPPC) neurons relate to large-scale brain activity during naturalistic video viewing in macaque monkeys. We found that neuron–BOLD functional maps exhibit strong context-dependent organization and capture contextual structure more robustly than neuronal spiking activity or fMRI frames alone. Despite this shared organization, neighboring dmPPC neurons often displayed markedly distinct whole-brain association patterns, revealing substantial local functional heterogeneity. These findings provide insight into how local neuronal populations participate in distributed cortical organization during naturalistic episodic processing.

## INTRODUCTION

The dorsomedial posterior parietal cortex (dmPPC) integrates sensory, cognitive, and motor information, playing an important role in episodic processing (Dadario & Sughrue, 2023; Suchan et al., 2002). Previous electrophysiological and fMRI studies have demonstrated substantial functional diversity within the dmPPC across multiple spatiotemporal scales (Margulies et al., 2009; Wang et al., 2024). Owing to its extensive anatomical and functional connectivity (Cavanna & Trimble, 2006; Zilles et al., 2003), dmPPC function likely depends not only on local neuronal activity but also on dynamic interactions with distributed cortical networks (Li et al., 2019; Thomas Yeo et al., 2011). However, existing methodologies provide only partial access to these processes.

Electrophysiological recordings capture neuronal activity with high temporal precision but are spatially restricted, whereas fMRI reveals large-scale network organization but lacks single-neuron resolution (Finn et al., 2023; Logothetis, 2008; Siegel et al., 2012; Wang et al., 2024). Consequently, how individual dmPPC neurons participate in distributed brain-wide functional organization remains poorly understood. Crucially, recent neurophysiological evidence suggests that dmPPC neurons do not merely encode isolated sensory features but exhibit mixed selectivity for high-dimensional and temporally evolving variables during naturalistic experience (Wang et al., 2024; Zuo et al., 2026). This coding complexity implies that distinct neuronal populations within the dmPPC may dynamically engage different large-scale cortical networks depending on ongoing contextual demands. We therefore hypothesized that dmPPC neurons would exhibit heterogeneous whole-brain neuron–BOLD association patterns reflecting their distinct tuning profiles during naturalistic video processing.

To bridge the gap between microscopic neuronal tuning and macroscopic network dynamics, we adopted an integrated multimodal approach combining single-unit electrophysiology with whole-brain fMRI. Previous studies have examined relationships between electrophysiological activity and BOLD signals (Kahn et al., 2013; Logothetis et al., 2001), but relatively few have integrated neuronal spiking and fMRI measurements to investigate how single-neuron activity relates to large-scale brain organization during cognitive processing (Cheng et al., 2023; Yang et al., 2026). Recent advances in single-unit fMRI mapping (Park et al., 2017) have provided a framework for linking neuronal activity to distributed whole-brain functional patterns during naturalistic viewing. In this approach, the activity time course of each neuron is systematically compared with voxel-wise fMRI BOLD signals across the brain, generating a whole-brain neuron–BOLD association map (“functional map”) for each recorded neuron based on Spearman correlation coefficients. Using this method, previous studies demonstrated that neighboring neurons with similar sensory selectivity can nonetheless exhibit markedly distinct whole-brain functional maps (Park et al., 2017; Russ et al., 2023), revealing substantial functional heterogeneity within local neuronal populations. These findings suggest that local neuronal diversity may reflect differential participation in distributed cortical networks during naturalistic processing (Hasson et al., 2008; Kauppi et al., 2010; Meer et al., 2020; Russ & Leopold, 2015; Ye et al., 2018).

Building on these methodological advances, we investigated functional diversity within the dmPPC during naturalistic video viewing. Specifically, we examined how neuron–BOLD functional maps vary across neuronal populations under shared and distinct contextual conditions, and compared the sensitivity of this approach with conventional analyses based on neuronal spiking activity or fMRI frames alone. We hypothesized that dmPPC neurons would exhibit context-dependent whole-brain association patterns reflecting their heterogeneous tuning properties and differential engagement with distributed cortical networks. We therefore tested whether dmPPC neurons exhibit stable or context-dependent large-scale association patterns during naturalistic viewing. By characterizing these neuron-specific functional maps, our study aimed to provide a systems-level perspective on how local dmPPC activity relates to large-scale brain organization during naturalistic processing.

## METHODS

### Experimental design overview

This study combined awake fMRI and in vivo extracellular electrophysiological recordings acquired from separate cohorts of macaque monkeys viewing identical naturalistic video stimuli. One cohort (*n* = 2) underwent fMRI scanning during video viewing, whereas a second cohort (*n* = 3) viewed the same videos during extracellular neuronal recordings. The stimulus set consisted of 18 naturalistic video clips, each lasting 30 s, presented identically across both experimental modalities.

### Subjects

Five rhesus macaques (*Macaca mulatta*) participated in this study. The first cohort consisted of two adult female monkeys, DP (6 years old, 9 kg) and VL (6 years old, 7 kg), which participated in the awake-fMRI experiments conducted at the Institute of Neuroscience, Newcastle University, UK. All procedures for this cohort were approved by the UK Home Office under the Animal Scientific Procedures Act (1986) and complied with the European Directive (2010/63/EU) for the protection of animals used in scientific research.

The second cohort consisted of three adult male monkeys: Mercury (MM; 6 years old, 9 kg), Jupiter (JM; 6 years old, 8 kg), and Galen (GM; 8 years old, 9.2 kg), which participated in the extracellular electrophysiological experiments conducted at the Nonhuman Primate Research Center, East China Normal University, China. Eye movements were additionally monitored in monkey Galen during the electrophysiological recordings. All procedures involving animal care, surgery, experimentation, and pre-/post-surgical care were approved by the Institutional Animal Care and Use Committee of East China Normal University (approval codes: M020150902 and M020150902-2018).

### Video stimuli and presentation

A free-viewing paradigm was used in both the fMRI and electrophysiological experiments. Video stimuli were collected from YouTube and edited using VideoStudio X8 into 720p clips at 25 frames/s. The videos depicted naturalistic scenes, including primate social behaviors, non-primate animals in natural environments, and landscape footage. For example, one clip showed a juvenile macaque interacting with other macaques during social play (more details can be here in Wang et al., 2024).

The final stimulus set consisted of 18 video clips, each lasting 30 s. In the fMRI experiments, the clips were divided into two sets, each containing nine clips. In the electrophysiological experiments, the 18 clips were organized into six video lists, each consisting of three clips (90 s total duration per list). Importantly, each recorded neuron was exposed to one—and only one—video list during a recording session. Each video list was repeated 30 times within a daily session, and each session was repeated across two recording days, yielding a total of 12 electrophysiological sessions. Water rewards were delivered at the onset of video presentation and during blank-screen intervals between video repetitions.

### *In* vivo extracellular data acquisition

Monkeys were implanted with a multichannel SC32 acquisition system (Gray Matter Research, LLC, USA) targeting the dorsomedial posterior parietal cortex (dmPPC). During electrophysiological recordings, monkeys were seated in a custom-designed Plexiglas chair with head restraint while freely viewing one of six video lists, each containing three 30-s video clips. Neuronal activity from monkeys MM and JM was recorded using glass-coated electrodes (SC32, Gray Matter Research, LLC, USA), whereas recordings from monkey GM were obtained using 24-channel single-shank tungsten microelectrodes (LMA, MicroProbes, USA). Detailed descriptions of surgical procedures, recording locations, behavioral paradigms, and experimental protocols have been reported previously (Wang et al., 2024). Across all recording sessions, we obtained neuronal activity from 285 active recording sites and isolated a total of 375 units for analysis. Of these, 204 were well-isolated single units obtained from individual recording sites (one unit per site). The remaining 171 units were isolated from 81 recording sites exhibiting multi-unit activity, yielding either two units (72 sites) or three units (9 sites) per recording site after spike sorting.

### Electrophysiology data preprocessing

Neural signals were initially high-pass filtered online at 1000 Hz during acquisition. Raw extracellular signals were subsequently bandpass filtered between 0.1 and 5500 Hz and digitized at 30 kHz. Spike sorting was performed offline using Offline Sorter (Plexon Inc., USA) based on waveform peak amplitude, principal component features, autocorrelation structure, and spike width. Units with a mean firing rate below 1 Hz across video presentations were excluded from subsequent analyses.

### Awake monkey fMRI data acquisition

During fMRI experiments, two monkeys (DP and VL) were scanned while awake with head stabilization provided by a customized distortion-free head-post implant (Ortiz-Rios et al., 2018). The animals viewed the same 18 naturalistic video clips used in the electrophysiological experiments, with each clip repeated 2–3 times per session. A 15-s dark interval separated successive clips. Across nine fMRI sessions, each monkey viewed every clip 4–6 times in total. To minimize jaw movement during image acquisition, juice rewards were delivered only at the end of each scanning session.

MRI data were acquired using a 4.7-T vertical magnet running ParaVision 5.1 (Bruker BioSpin GmbH, Ettlingen, Germany). High-resolution anatomical images were first collected using an MP-RAGE sequence (0.61 × 0.61 × 0.61 mm³ resolution; TE = 3.74 ms; TR = 2000 ms; flip angle = 90°). Functional images were then acquired using gradient-echo echo-planar imaging (EPI) (1.2 × 1.2 × 1.2 mm³ resolution; TE = 21 ms; TR = 1500 ms; flip angle = 65°) with a 4-channel phased-array coil (https://www.wkscientific.com).

### fMRI data preprocessing

All fMRI preprocessing was performed using the AFNI and SUMA software packages (Cox, 1996). Preprocessing steps included slice-timing correction, motion correction, anatomical co-registration, spatial smoothing, global signal scaling, and nuisance regression. Functional data from each session were aligned to the NIMH Macaque Template (NMT v2; Jung et al., 2021) to facilitate cross-subject visualization and group-level analysis. For each session, BOLD time series corresponding to repeated presentations of the same video clip were extracted separately for all 18 clips. The repeated trials were then averaged to generate a mean stimulus-evoked time series for each video clip.

### Construction of neuron-BOLD functional maps

To account for differences in temporal resolution between electrophysiological and fMRI recordings, neuronal spiking time series were first downsampled to match the fMRI sampling rate. The resulting spike-rate time series were then convolved with a canonical hemodynamic response function (gamma function; Figure 1A). For each neuron, the convolved time series was concatenated across the three 30-s video clips within a given video list, yielding a 90-s stimulus-locked neuronal time course.

**Figure 1.**
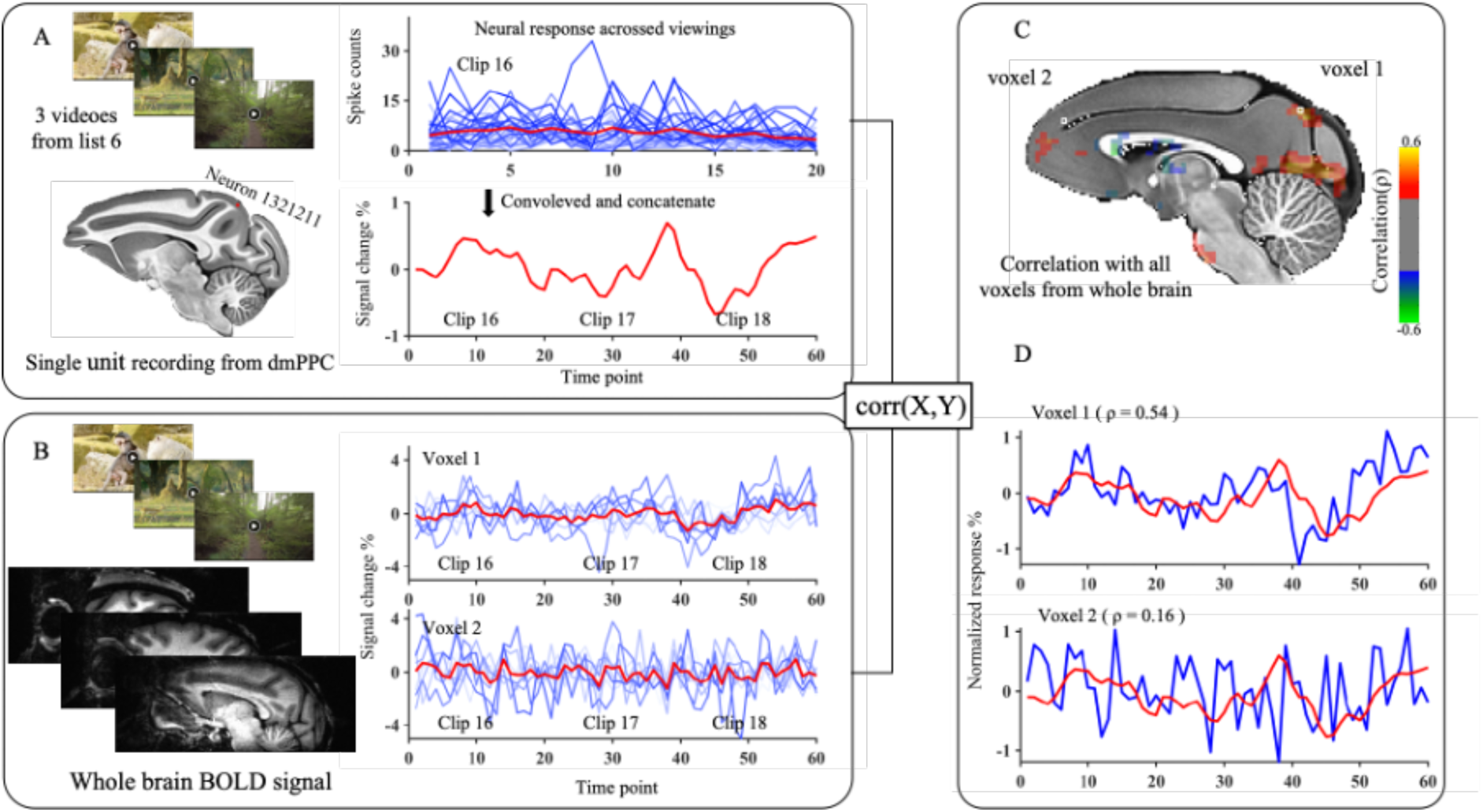
Workflow of neuron–BOLD functional map construction. Illustration of the procedure used to construct a whole-brain functional map for a single neuron. **(A)** Convolved and concatenated spiking-rate profile of an example neuron during presentation of a video list containing three 30-s clips. **(B)** Example voxel-wise fMRI BOLD signals recorded during presentation of the same video sequence. **(C)** Whole-brain functional map of the example neuron, generated by computing the correlation between the neuron’s convolved spiking profile (A) and voxel-wise fMRI BOLD signals across the brain. **(D)** Example comparisons between the neuron’s convolved spiking profile and two voxel BOLD signals, illustrating how temporal correspondence between neuronal and fMRI signals gives rise to the neuron–BOLD association used to construct the functional map. See also **Figure S1** a schematic overview illustrating the full methodological pipeline.

Correspondingly, voxel-wise BOLD time series were extracted and concatenated across the same video sequence (Figure 1B). Functional maps were subsequently generated by computing Spearman’s rank correlation coefficients between each neuron’s convolved time series and the time series of every voxel across the brain (Figure 1C). This procedure produced a whole-brain functional map for each neuron, reflecting the degree to which voxel-wise BOLD signals covaried with that neuron’s activity during naturalistic viewing. By leveraging identical video stimuli and synchronized stimulus timing across electrophysiological and fMRI experiments, this approach enabled direct integration of neuronal and whole-brain imaging data to generate neuron–BOLD functional maps for individual neurons (Figure 1D). Importantly, each neuron was recorded during the presentation of one—and only one—of the six video lists. Consequently, each neuron’s functional map was intrinsically linked to a specific 90-s contextual sequence composed of three consecutive 30-s clips, and neurons recorded within the same session therefore shared the same contextual condition. To further improve the clarity of the analytical workflow and conceptual interpretation of the neuron–BOLD functional maps, we have also added a schematic overview figure illustrating the full pipeline from naturalistic video viewing and neuronal recordings to construction and comparison of stimulus-locked neuron–BOLD association maps (Figure S1).

Statistical reliability of neuron–BOLD correlation maps was evaluated using a voxel-wise non-parametric stationary bootstrap procedure adapted from Park et al. (2017). Voxels exhibiting reliable neuron–BOLD association values across the population were retained for subsequent analyses.

### Selection of a subset of voxels

To restrict analyses to voxels exhibiting reliable stimulus-evoked activity, we generated an amplitude mask based on the root mean square (RMS) of each voxel’s preprocessed BOLD time series. Voxels with RMS amplitudes below 0.37% signal change (corresponding to the 50th percentile of the voxel-wise amplitude distribution; Figure S2A) were excluded from subsequent analyses. This procedure follows approaches used in previous single-unit fMRI mapping studies (Park et al., 2017) and reduces the contribution of voxels with minimal stimulus-related fluctuations. For population-level analyses, neuron–BOLD functional maps were further restricted to voxels exhibiting robust association values across the neuronal population. Specifically, only voxels showing absolute correlation values greater than 0.35 with at least 5% of recorded neurons were retained. After thresholding, each neuron’s functional map was represented as a vector containing 5,027 voxel-wise association values. The resulting amplitude maps and final voxel masks are shown in Figure S2.

### Definition and selection of critical fMRI frames

Critical fMRI frames (*n* = 486) were defined as the top 15% of fMRI frames exhibiting the highest BOLD signal amplitude within the dmPPC region of interest across the entire video-viewing dataset, following the approach of Liu and Duyn (2013). The dmPPC region was defined using area PEc from the CHARM atlas (Cortical Hierarchy Atlas of the Rhesus Macaque, Jung et al., 2021).

For each fMRI frame, the mean BOLD signal amplitude within the dmPPC ROI was calculated, and frames exceeding the 85th percentile of the amplitude distribution were selected for subsequent analyses. These high-amplitude frames were used to characterize context-dependent fMRI patterns and to compare frame-based representations with neuron–BOLD functional maps and neuronal spiking activity.

### Comparison of context-dependent organization across modalities

To characterize dmPPC function across different analytical scales, we compared three complementary modalities: (1) neuron–BOLD functional maps, representing whole-brain association patterns between individual dmPPC neurons and voxel-wise BOLD signals; (2) neuronal spiking activity, representing local electrophysiological dynamics; and (3) critical fMRI frames, representing high-amplitude dmPPC-related whole-brain fMRI activity patterns during video viewing.

To determine whether neuron–BOLD functional maps captured contextual organization more robustly than neuronal spiking activity or critical fMRI frames alone, we applied a unified similarity-based analytical framework across all three modalities. For each neuron (or each critical fMRI frame), all comparisons were divided into two conditions: a within-video condition, consisting of neurons or frames associated with the same video list, and an across-video condition, consisting of neurons or frames associated with different video lists. For neuron–BOLD functional maps, similarity values were calculated using Pearson correlation coefficients between the vectorized functional map of a given neuron and those of all other neurons. Correlation coefficients were Fisher z-transformed to stabilize variance, and absolute transformed values (|z|) were used as similarity metrics. Welch’s *t*-tests were then performed separately for each neuron to compare within-video and across-video similarity distributions. Resulting *p*-values were corrected for multiple comparisons using false discovery rate (FDR) correction, and Cohen’s *d* was calculated to estimate effect size. Equivalent analyses were performed for neuronal spiking activity and critical fMRI frames using the corresponding modality-specific representations.

At the population level, similarity distributions across modalities and comparison conditions were further evaluated using a Scheirer–Ray–Hare test, a non-parametric alternative to two-way ANOVA appropriate for unbalanced sample sizes and non-normal similarity distributions. Similarity values were first rank-transformed, and the effects of modality (functional maps, spiking activity, or critical fMRI frames), comparison type (within-video vs. across-video), and their interaction were subsequently assessed. Low-dimensional visualization of functional maps, neuronal spiking activity, and critical fMRI frames was performed using Uniform Manifold Approximation and Projection (UMAP, McInnes et al., 2018).

### Clustering analysis of neuron–BOLD functional maps

K-means clustering was applied in two separate analyses of neuron–BOLD functional maps. First, clustering was used to determine whether neighboring neurons recorded simultaneously from the same electrode exhibited similar or divergent functional map organization. Second, clustering was used to examine functional heterogeneity among neurons recorded under the same video condition. For both analyses, vectorized neuron–BOLD functional maps were used as input features for standard k-means clustering. The optimal number of clusters (k) was determined using silhouette analysis, which quantified clustering quality across different values of k. The cluster number yielding the highest silhouette score was selected for subsequent analyses.

## RESULTS

### Neuron-BOLD functional maps exhibit stronger context-dependent organization than spiking activity or critical fMRI frames alone

To evaluate whether neuron–BOLD functional maps capture context-dependent organization more robustly than conventional electrophysiological or fMRI-based measures, we compared similarity structures across three modalities derived from dmPPC activity (Figure 2A): neuron–BOLD functional maps (Figure 2B), neuronal spiking patterns (Figure 2C), and critical fMRI frames (Figure 2D). We first quantified similarity relationships between within-video and across-video pairs across the three modalities. Representational similarity matrices revealed pronounced block-like structure in neuron–BOLD functional maps, indicating greater similarity among neurons recorded during presentation of the same video sequence compared with neurons associated with different video sequences (Figure 3A). In contrast, this context-dependent organization was substantially weaker for neuronal spiking patterns (Figure 3B) and critical fMRI frames (Figure 3C).

**Figure 2.**
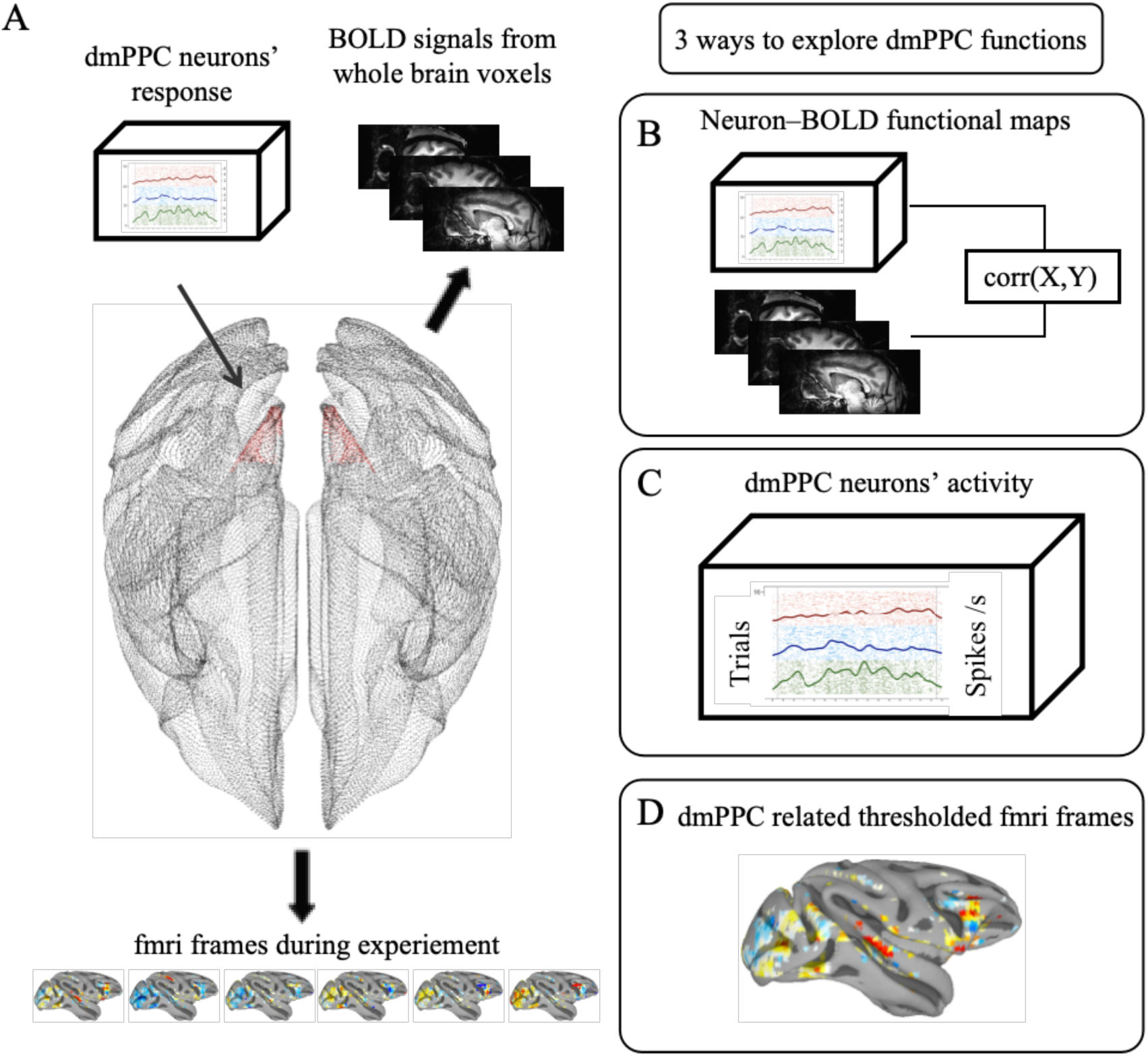
Three complementary approaches for characterizing dmPPC function under viewing task. **(A)** Schematic overview of the multimodal framework used to investigate dmPPC function during naturalistic video viewing. Simultaneously aligned neuronal spiking responses and whole-brain fMRI signals were analyzed using three complementary representations: **(B)** neuron–BOLD functional maps, generated by correlating each neuron’s convolved spiking profile with voxel-wise fMRI BOLD signals across the brain; **(C)** neuronal spiking activity patterns derived directly from dmPPC neurons; and **(D)** thresholded critical fMRI frames representing high-activity whole-brain fMRI states associated with the dmPPC region.

**Figure 3.**
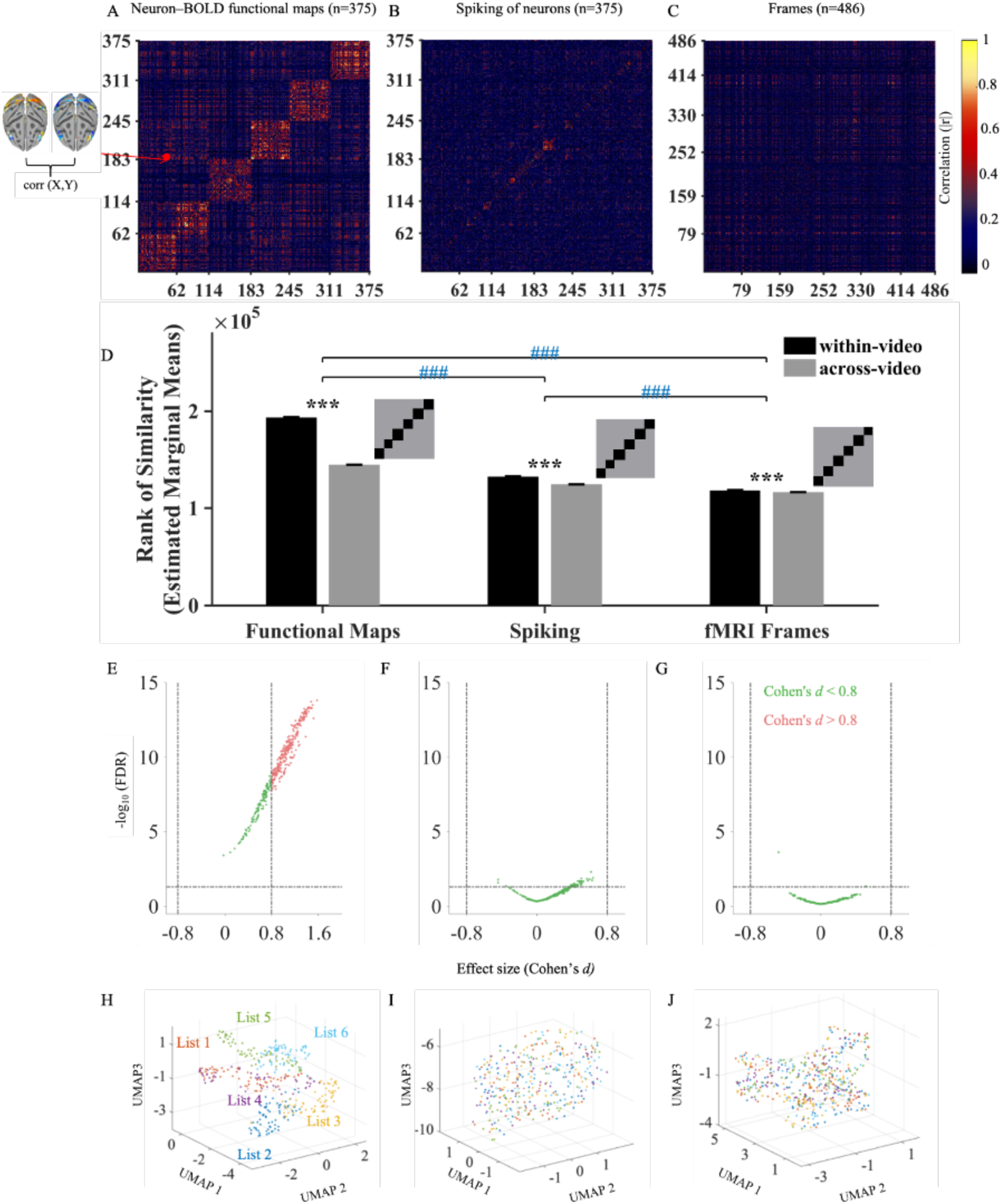
Context-dependent similarity structure revealed by functional maps, neuronal spiking patterns, and critical fMRI frames. **(A–C)** Modality-specific similarity matrices showing the absolute correlation between pairs of representations derived from neuron–BOLD functional maps (A), neuronal spiking patterns (B), and critical fMRI frames (C). In **(A)**, each matrix entry represents the spatial correlation between the whole-brain functional maps of two neurons. Functional maps were generated by correlating each neuron’s convolved spiking profile with voxel-wise fMRI BOLD signals during presentation of the same (matching) video sequence, thereby capturing stimulus-locked neuron–BOLD association profiles. For visualization purposes, neurons are ordered according to the video sequence during which they were recorded, producing the block structure visible in the matrix. The schematic on the left illustrates the computation of a single matrix entry, corresponding to the absolute correlation between the functional maps of neuron *i* and neuron *j*. In **(B)** and **(C)**, the same similarity procedure was applied to neuronal spiking patterns and critical fMRI frames, respectively. For visualization purposes, all diagonal values (self-correlations of 1.0) were set to 0 to prevent them from obscuring off-diagonal differences in the color scale. **(D)** Comparison of context-dependent similarity effects across modalities. Bars show estimated marginal means of rank-transformed similarity values derived from a non-parametric two-way ANOVA (Scheirer–Ray–Hare test). Black bars indicate within-video similarity, and grey bars indicate across-video similarity. Error bars denote standard error of means across pair-wise correlation values. Asterisks (***) denote significant within-vs. across-video differences within each modality (Bonferroni-corrected p < 0.001). Blue hash marks (###) indicate significant interaction contrasts between modalities (Bonferroni-corrected p < 0.001). Insets illustrate hypothetical within-video (diagonal blocks) versus across-video (off-diagonal) similarity structure. **(E–G)** Volcano plots showing modality-specific similarity differences at the neuron/frame level for functional maps (E), neuronal spiking patterns (F), and critical fMRI frames (G). The x-axis shows effect size (Cohen’s *d*), and the y-axis shows statistical significance (−log10 FDR-adjusted *p*-value). Dashed lines indicate thresholds for significance (FDR *p* = 0.05) and large effect size (|*d*| = 0.8). Points meeting both criteria are highlighted in red. **(H–J)** UMAP projections illustrate clustering by video condition for functional maps (H), but not for neuronal spiking patterns (I) or critical fMRI frames (J). Color coding in (I) and (J) correspond to the same video conditions shown in (H).

To directly compare the magnitude of the context-dependent effect across modalities, we performed a non-parametric two-way ANOVA (Scheirer–Ray–Hare test) using modality and comparison type (within-video vs. across-video) as factors. This analysis revealed significant main effects of modality and comparison type, as well as a significant interaction between the two factors, indicating that the strength of the context-dependent effect differed across modalities. Post-hoc comparisons showed that neuron–BOLD functional maps exhibited significantly larger within-video versus across-video differences than both neuronal spiking patterns and critical fMRI frames, whereas spiking patterns also showed a significantly larger effect than critical fMRI frames (all Bonferroni-corrected *p* < 0.001; Figure 3D). Together, these findings indicate that neuron–BOLD functional maps preserve context-dependent structure more strongly than either neuronal spiking activity or critical fMRI frames alone.

To further examine these effects at the individual neuron/frame level, we computed effect sizes (Cohen’s *d*) and statistical significance for within-video versus across-video similarity differences across all modalities. Neuron–BOLD functional maps showed widespread large effect sizes (*d* > 0.8) together with significant group differences (FDR-corrected *p* < 0.05; Figure 3E), whereas neuronal spiking patterns (Figure 3F) and critical fMRI frames (Figure 3G) exhibited substantially weaker effects and did not reach the same effect-size threshold. Finally, we visualized the similarity structure of the three modalities using UMAP dimensionality reduction. Neuron–BOLD functional maps formed distinct clusters corresponding to video context (Figure 3H), whereas neuronal spiking patterns (Figure 3I) and critical fMRI frames (Figure 3J) showed substantially weaker clustering. These results further support the conclusion that neuron–BOLD functional maps capture context-dependent organization more robustly than either neuronal spiking activity or critical fMRI frames alone.

### Classification of clip-specific functional maps across contextual conditions

As shown in the previous section, neuron–BOLD functional maps exhibited significantly greater similarity within video-list conditions than across different video lists, indicating strong context-dependent organization of dmPPC neuron–BOLD association patterns. We next asked whether the same neuron would exhibit distinct whole-brain association patterns across different video clips, and whether these clip-specific functional maps could be reliably classified according to contextual condition.

Each recorded neuron was associated with a video list containing three distinct 30-s video clips. By separating the activity corresponding to each clip, we generated three clip-specific functional maps for each neuron, allowing us to examine how the same neuron’s whole-brain association structure varied across different contextual conditions. To minimize the influence of spurious correlations arising from the shorter 30-s analysis windows, all analyses were restricted to the pre-defined voxel mask derived from the full 90-s recordings using the stationary bootstrap procedure described above.

Visual inspection revealed substantial context-dependent variation in neuron–BOLD association patterns across clips. Figure 4A shows example clip-specific functional maps from 12 neurons recorded during presentation of clips 1–3 and clips 4–6. Even within the same neuron, different clips produced markedly distinct whole-brain association patterns, demonstrating that neuron–BOLD organization dynamically changes with contextual condition. At the same time, clip-specific functional maps derived from different neurons showed relatively consistent large-scale organization within the same clip condition. For example, during clip 1, many neurons exhibited positive associations with ventrolateral prefrontal, lateral motor, auditory belt, somatosensory, and visual cortical regions, together with negative associations in ventral premotor cortex. Distinct but internally consistent spatial patterns were similarly observed during clips 2 and 3 (Figure 4A). These findings indicate that dmPPC neurons exhibit both shared context-dependent organization and substantial functional heterogeneity across contextual conditions.

**Figure 4.**
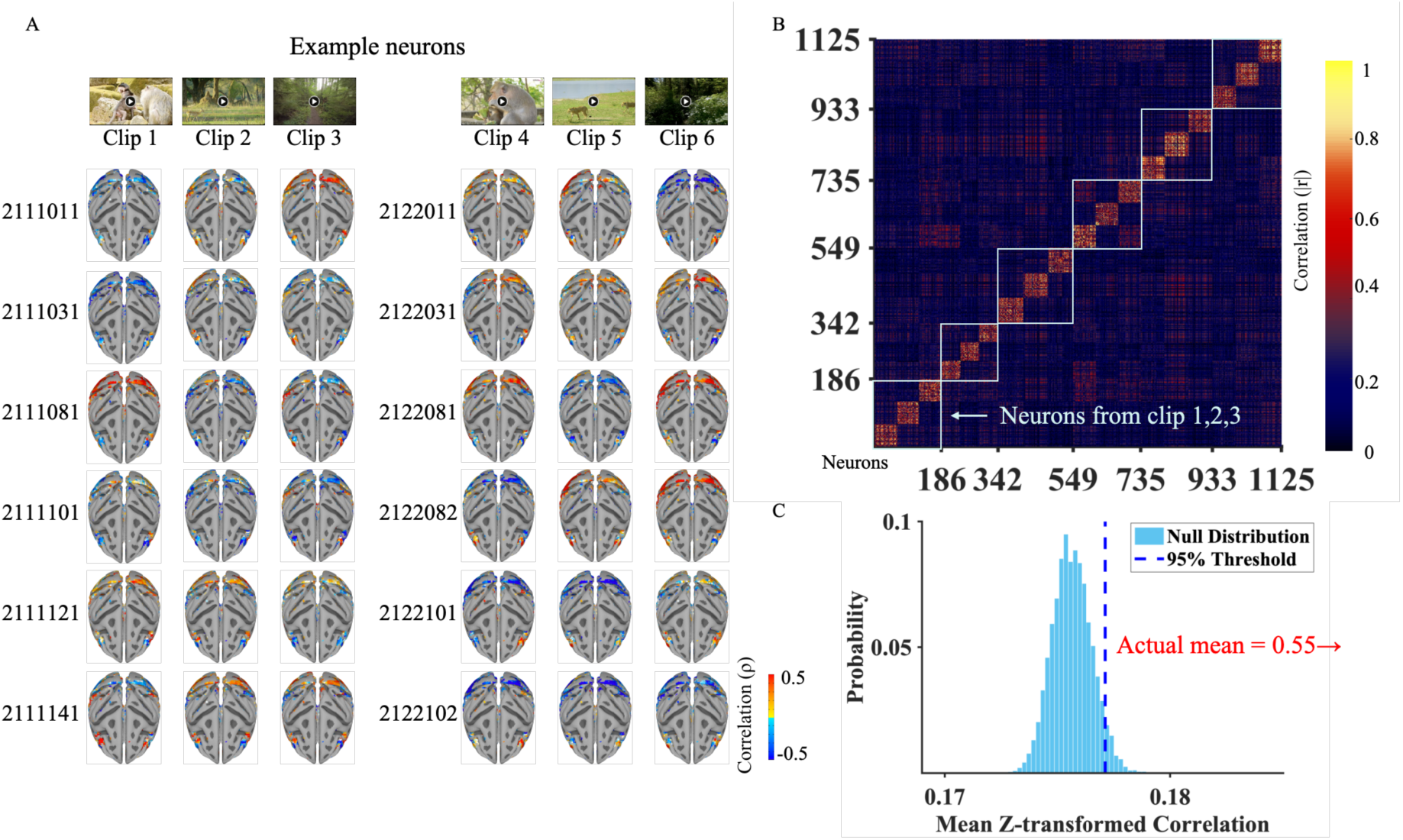
Context-dependent similarity structure of neuron–BOLD functional maps across video clips. **(A)** Example neuron–BOLD functional maps obtained during presentation of six different video clips. Each row represents a different neuron, and each column corresponds to a specific video clip condition. Distinct spatial patterns of neuron–BOLD association are observed across clips. **(B)** Representational similarity matrix (RSM) comparing functional maps across all neurons and video clips. Each matrix entry represents the absolute correlation between a pair of neuron–BOLD functional maps computed within a single clip condition. Functional maps are grouped according to video clip identity, producing the block structure visible along the diagonal. Functional maps corresponding to clips 1–3 from panel (A) are indicated in the lower-left region of the matrix. Clip 1 depicts social interactions between macaques; clip 2 shows a herd of deer moving through grassland; and clip 3 shows movement along a forest pathway. **(C)** Permutation analysis evaluating within-clip similarity. The blue histogram represents the null distribution generated from 10,000 permutations. The red arrow indicates the observed mean within-clip similarity, which exceeded the 95% confidence interval of the null distribution (*p* < 0.0001), demonstrating significantly higher similarity among functional maps derived from the same video context.

To quantitatively evaluate whether clip-specific functional maps could be reliably classified according to contextual condition, we performed a permutation-based similarity analysis. We first computed absolute Pearson correlation coefficients between all pairs of the 1,125 clip-specific functional maps, generating a 1,125 × 1,125 representational similarity matrix (RSM; Figure 4B). The resulting matrix exhibited a prominent block structure, indicating that functional maps derived from the same video clip were more similar to one another than to maps derived from different clips. To assess statistical significance, we constructed a null distribution by randomly shuffling clip labels across maps (10,000 iterations). For each iteration, we identified pseudo-within-clip pairs based on shuffled labels and computed their mean similarity value. The observed within-clip similarity substantially exceeded the null distribution and lay beyond the 95th percentile threshold (Figure 4C). None of the 10,000 shuffled iterations equaled or exceeded the observed value (*p* < 0.0001), demonstrating that clip-specific functional maps contain reliable, non-random contextual structure.

### Reliability of short-duration stimulus-evoked BOLD responses

To evaluate whether the relatively short (30-s) video clips provided sufficiently reliable fMRI signals for neuron–BOLD association analysis, we performed split-half reliability analyses across repeated presentations of each clip (Figure S3A). Repeated trials were randomly divided into two independent groups, and voxel-wise correlations were computed between the mean BOLD time courses of the two groups. Reliable stimulus-locked responses were observed across widespread cortical regions, particularly within visual and parietal areas. To further assess statistical significance, we compared the observed split-half reliability against null distributions generated by temporal-shuffling permutation tests (1,000 iterations; Figure S3B). For the majority of video clips, observed reliability values exceeded the null distributions, indicating that the measured BOLD responses reflected structured stimulus-driven activity rather than random fluctuations or slow hemodynamic drift. Together, these analyses demonstrate that the 30-s clips were sufficient to elicit reproducible context-dependent BOLD responses suitable for subsequent neuron–BOLD association analyses.

### Neighboring dmPPC neurons exhibit both local similarity and functional heterogeneity

At the population level, neurons recorded during the same video context formed distinct functional clusters (Figure 4), indicating a dominant macro-scale organization in which shared contextual conditions drive global similarity in neuron–BOLD association structure. To further examine the fine-grained organization of these patterns, we investigated whether neighboring neurons recorded simultaneously from the same electrode—and therefore sharing the same local environment and stimulus context—would exhibit similar or divergent whole-brain association patterns. Previous studies have shown that neighboring neurons can exhibit similar functional properties because of shared local circuit organization (Hubel & Wiesel, 1962; Smith & Kohn, 2008). However, neighboring neurons in higher-order association cortex may also display distinct large-scale functional relationships, as previously demonstrated in inferotemporal cortex (Park et al., 2017).

We found evidence for both local similarity and substantial local heterogeneity among neighboring dmPPC neurons. Two example neurons recorded simultaneously from the same electrode illustrate this divergence (Figure 5A-B). Neuron #2321191 exhibited negative associations with visual and superior temporal regions (ρ = −0.40; Figure 5A), whereas the neighboring neuron #2321192 showed widespread positive associations with visual cortical areas (ρ = 0.43; Figure 5B). Similar heterogeneity was observed across multiple simultaneously recorded neuronal pairs (Figure 5C), although many neighboring neurons also displayed broadly similar association patterns, consistent with shared stimulus-driven organization during video viewing.

**Figure 5.**
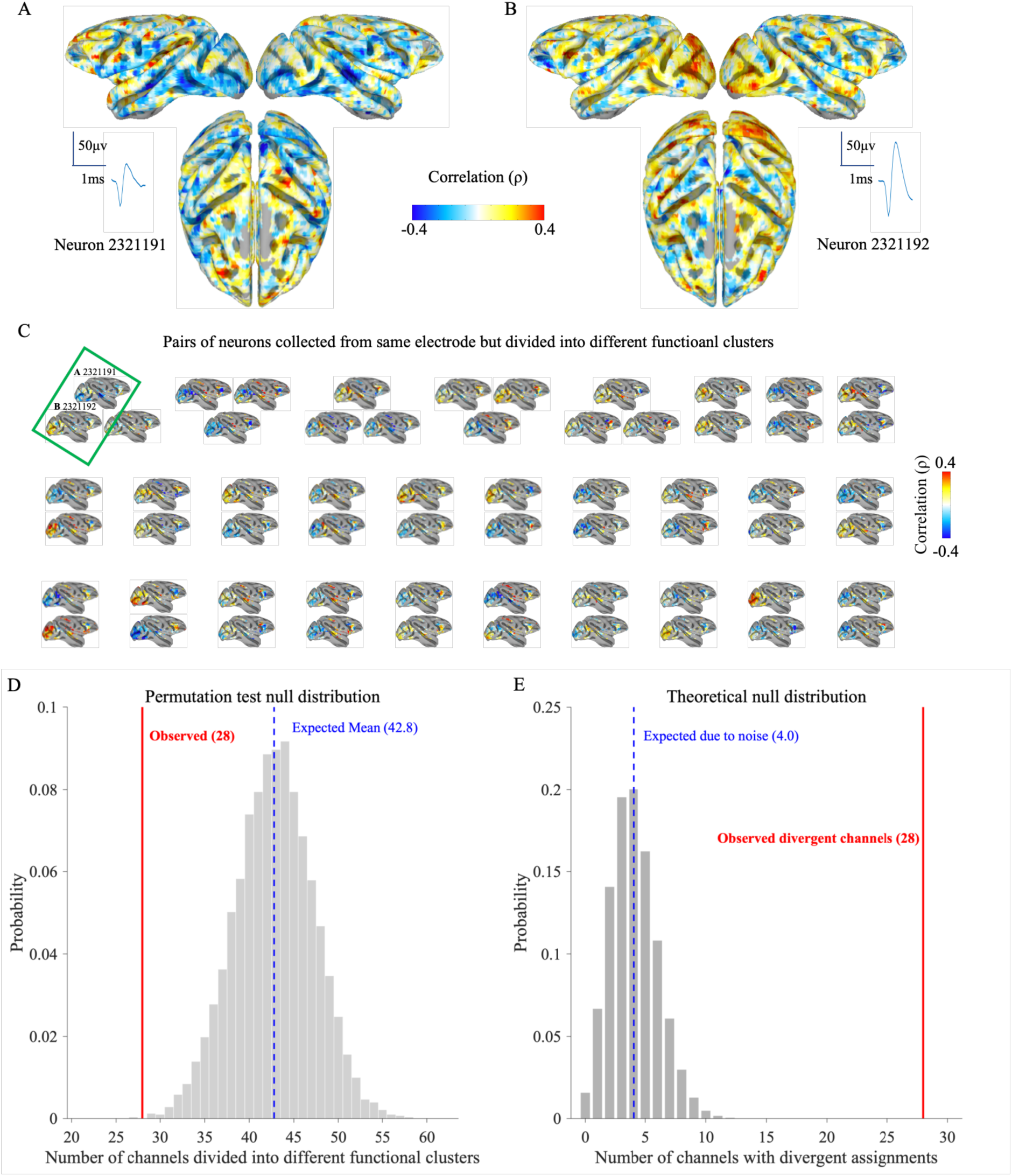
Local functional heterogeneity among simultaneously recorded dmPPC neurons. **(A–B)** Example whole-brain neuron–BOLD functional maps for neurons #2321191 (A) and #2321192 (B), simultaneously recorded from the same electrode in monkey JM. Despite being recorded from the same local site, the two neurons exhibit distinct neuron–BOLD association patterns. Insets show example spike waveforms for each neuron. **(C)** Functional maps from neurons simultaneously recorded on the same electrode but assigned to different functional clusters. Each panel represents neurons recorded from one electrode channel. The green rectangle highlights the example neurons shown in panels (A) and (B). The third neuron from the same channel (#2321193), not enclosed by the rectangle, exhibits a functional map more similar to neuron #2321192 than to neuron #2321191. Across the dataset, 28 of 80 multi-neuron recording channels contained neurons assigned to different functional clusters. **(D)** Permutation analysis evaluating spatial clustering of functional profiles. Gray histogram shows the null distribution generated by randomly shuffling cluster labels across neurons (10,000 permutations). The observed number of mixed-cluster channels (red line) was significantly lower than the null expectation (blue dashed line; *p* = 0.008), indicating that neighboring neurons are generally more likely to share similar functional profiles than expected by chance. **(E)** Assessment of local functional heterogeneity using an exact binomial test assuming a 5% probability of divergent cluster assignment due to system noise. The observed number of divergent channels (red line) significantly exceeded the expected noise-derived level (blue dashed line; *p* < 0.0001), demonstrating that local functional heterogeneity cannot be explained solely by noise.

To systematically quantify these effects, we analyzed 81 of the 285 electrodes (28.4%) that simultaneously recorded two or three isolated neurons (9 electrodes yielded three neurons; 72 electrodes yielded two neurons). We applied k-means clustering (*k* = 2) to the neuron–BOLD functional maps generated from the 90-s video sequences and examined whether neighboring neurons were assigned to the same or different clusters. Among the 81 electrodes, 28 (34.6%) contained neurons assigned to different functional clusters (Figure 5C). We first tested whether neighboring neurons were generally more similar to each other than expected by chance using permutation analysis (10,000 iterations). If cluster assignments were completely independent of spatial proximity, the expected number of mixed-cluster electrodes would be 42.8. The observed number of mixed-cluster electrodes (28) was significantly lower than this null expectation (*p* = 0.008), indicating that neighboring neurons are generally more likely to share similar functional profiles than expected by chance (Figure 5D).

We next asked whether the observed heterogeneity could nonetheless be explained solely by system noise. Under a conservative null hypothesis assuming complete local homogeneity with a 5% system error rate, the expected number of mixed-cluster electrodes was only 4.05 out of 81. In contrast, we observed 28 mixed-cluster electrodes. An exact binomial test demonstrated that this level of local heterogeneity was highly unlikely to arise from noise alone (*p* < 0.0001; Figure 5E). Together, these findings indicate that neighboring dmPPC neurons exhibit a nested organizational structure: shared contextual conditions produce broad local similarity in neuron–BOLD association patterns, while substantial functional heterogeneity persists within local microcircuits.

### Context-dependent neuron–BOLD association patterns across dmPPC-related cortical regions

While the preceding analyses characterized neuron–BOLD association patterns at the voxel level, we next examined how these patterns were organized across distributed cortical and subcortical regions associated with dmPPC function. Regions of interest (ROIs) were defined using the CHARM atlas (Cortical Hierarchy Atlas of the Rhesus Macaque, Jung et al., 2021) and SARM atlas (Subcortical Atlas of the Rhesus Macaque, Hartig et al., 2021), including areas PEc and PEci (representing dmPPC), primary motor cortex (M1), posterior cingulate gyrus (PCgG), parahippocampal cortex (paraHi), rhinal cortex (Rh), medial orbitofrontal cortex (mOFC/vmPFC), dorsolateral prefrontal cortex (dlPFC), hippocampus (Hi), and visual cortical areas V1–V3 (Figure 6A). We asked whether neuron–BOLD association patterns were uniformly distributed across these regions or instead exhibited region- and context-dependent organization during video viewing (Cavanna & Trimble, 2006; S. Zhang & Li, 2012).

**Figure 6.**
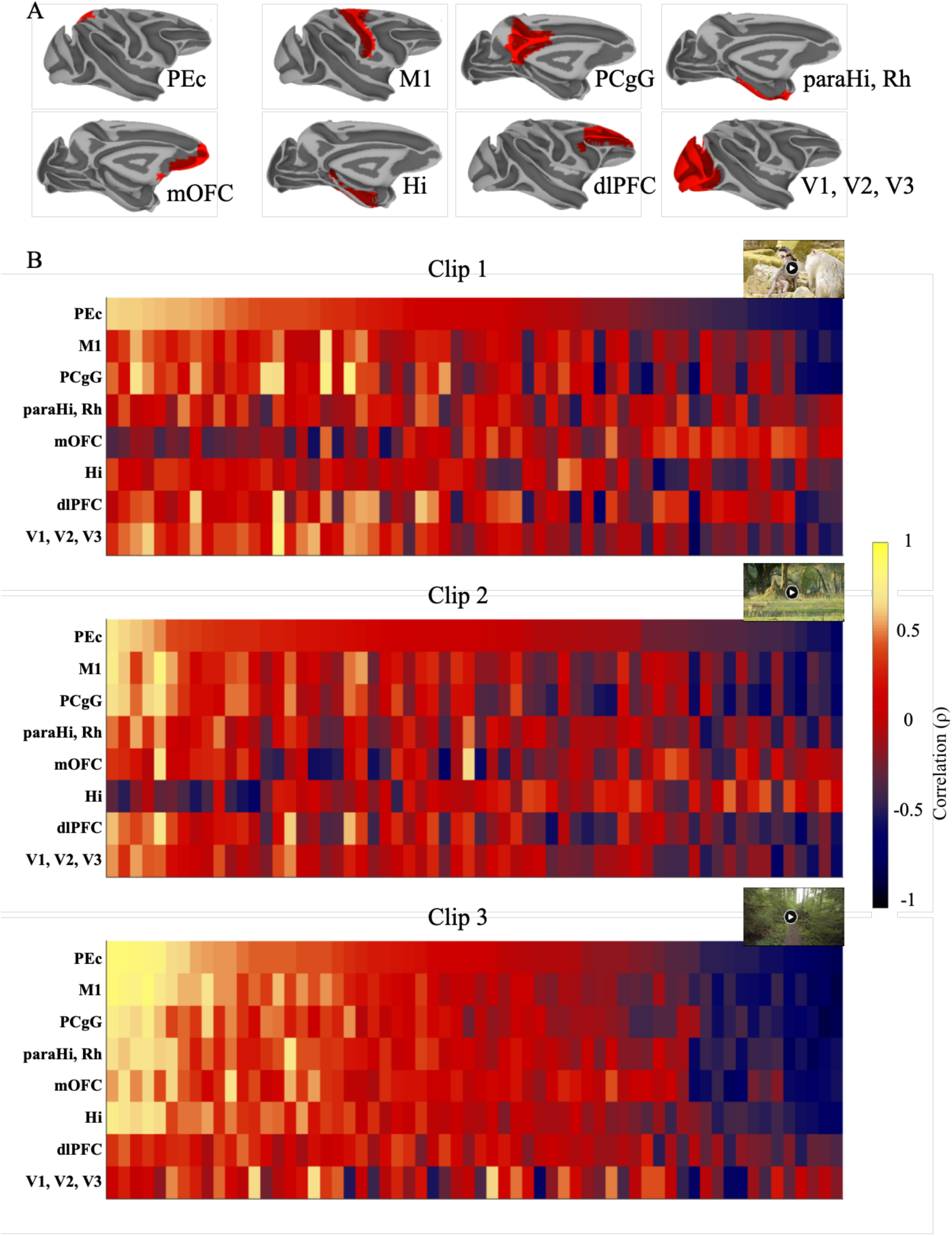
Context-dependent neuron–BOLD association patterns across cortical regions-of-interest (ROI). **(A)** Masks of dmPPC-related ROIs. Abbreviations: PEc, areas PEc and PEci (representing dmPPC); M1, primary motor cortex; PCgG, posterior cingulate gyrus; paraHi, parahippocampal cortex; Rh, rhinal cortex; mOFC, medial orbitofrontal cortex (vmPFC); Hi, hippocampus; dlPFC, dorsolateral prefrontal cortex; V1, primary visual cortex; V2, visual area 2; V3, visual area 3. **(B)** Heatmaps showing neuron–BOLD association values across ROIs during viewing of video clips 1–3. Each matrix corresponds to one video clip condition. Columns represent neurons and rows represent ROIs. Within each matrix, neurons are sorted in descending order according to their association strength within the dmPPC (PEc; first row), computed as the Spearman correlation between each neuron’s convolved spiking time series and ROI-averaged fMRI BOLD signals. This ordering reveals both shared and divergent neuron–BOLD association patterns across cortical regions and video contexts. Corresponding results for the remaining 15 video clips are shown in Figure S4.

Neuron–BOLD association values were computed as Spearman correlations between each neuron’s convolved spiking profile and ROI-averaged fMRI BOLD signals for all 1,125 clip-specific functional maps across the 18 video clips. Several ROIs exhibited highly structured association patterns. In particular, M1 and PCgG displayed consistent association gradients aligned with neuronal ordering in the dmPPC (PEc; Figure 6B). This gradient-like organization was robustly preserved across all 18 video clip conditions (clips 4 to 18 shown in Figure S4).

In contrast, higher-order cognitive regions exhibited substantially more heterogeneous and context-dependent association patterns. Prefrontal and medial temporal regions diverged markedly from the stable gradients observed in sensorimotor and cingulate regions, with association structure varying across video contexts. For example, during clips 1 and 6, the mOFC exhibited association patterns that were inversely related to the PEc gradient. Similarly, hippocampal association patterns were opposite to the PEc gradient specifically during clips 2, 6, and 18 (Figure 6B; Figure S4). These findings indicate that neuron–BOLD association structure within higher-order cortical systems dynamically changes across contextual conditions rather than remaining spatially fixed.

To determine whether these effects simply reflected broad brain-wide association levels, we compared within-ROI association values against the mean association values across voxels outside the ROI. For each of the 18 video clips, we calculated the difference between the mean within-ROI association and the extra-ROI mean association (Figure S5). The context-dependent patterns observed in higher-order cognitive regions remained robust after this control analysis, including the clip-specific inversions observed in mOFC and hippocampal regions. Together, these findings indicate that dmPPC neuron–BOLD association patterns are not uniformly distributed across the brain, nor are they reducible to nonspecific global association structure. Instead, dmPPC neurons exhibit dynamic and context-dependent coupling with distributed cortical and medial temporal systems during naturalistic video processing.

### Distinct functional clusters emerge within shared contextual conditions

We further observed that even within a single video clip condition, neuron–BOLD functional maps exhibited substantial functional heterogeneity. This heterogeneity was particularly evident in association patterns involving prefrontal and visual cortical regions (Figure 4A), suggesting that shared contextual conditions do not produce uniform large-scale organization across dmPPC neurons. To quantify this variability, we performed clustering analysis separately for the clip-specific functional maps derived from each video clip. Correlation coefficients from the same subset of valid voxels defined using the corresponding 90-s video list were used as input features for k-means clustering. The optimal number of functional clusters for each clip was determined using silhouette analysis (Figure S6). Depending on the video clip, neurons segregated into either two clusters (clips 1, 2, 3, 7, 8, 9, 10, 12, 13, 14, 15, 16, and 18), three clusters (clips 4, 5, and 17), four clusters (clip 6), or six clusters (clip 11) (Figure 7). Across all 18 clips, this analysis identified a total of 45 functional clusters.

**Figure 7.**
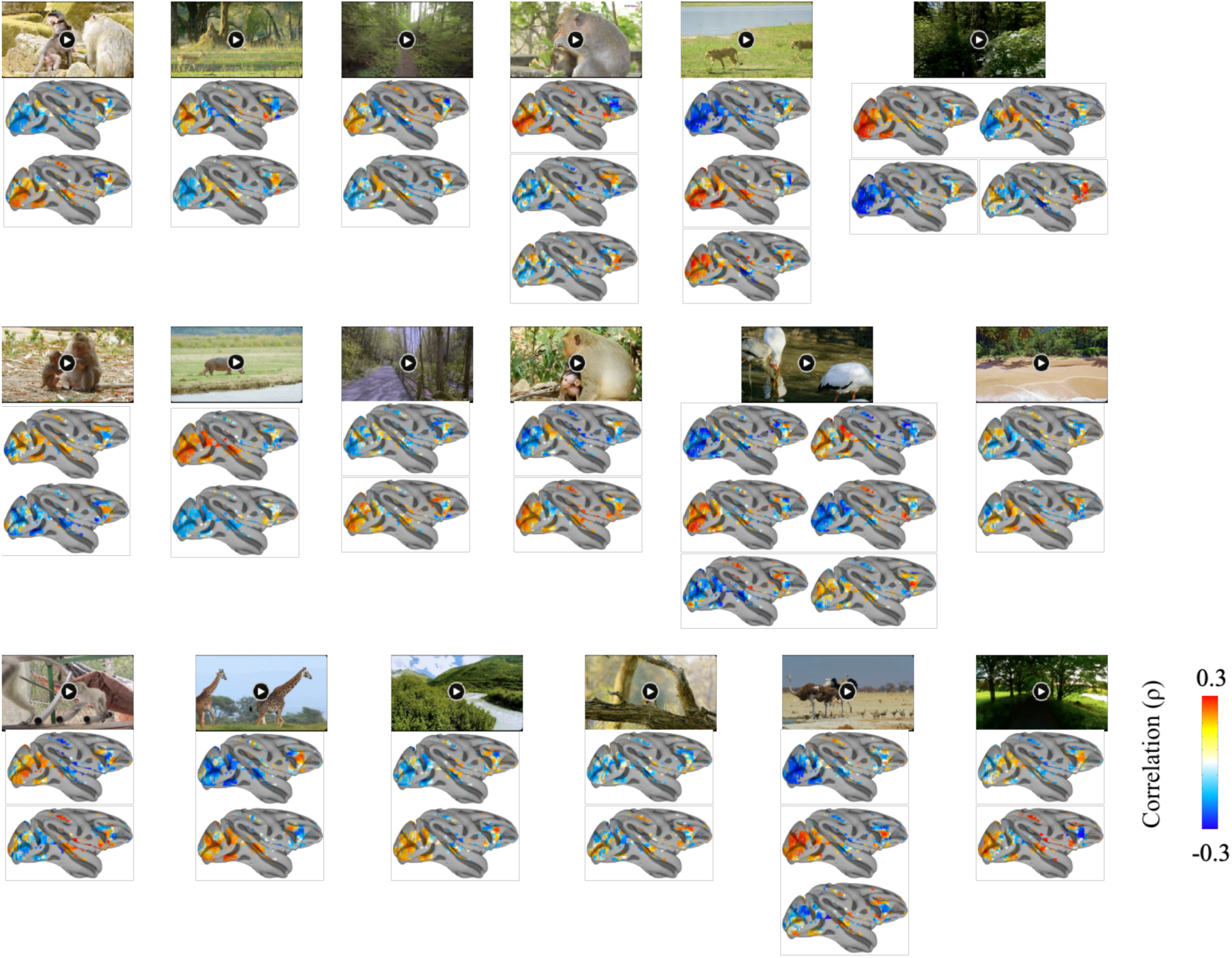
Functional clustering of dmPPC neuron–BOLD maps across video contexts. For each of the 18 video clips, neuron–BOLD functional maps were grouped using k-means clustering based on similarity of whole-brain association patterns. Each panel corresponds to one video clip condition, and each brain map represents the average functional map of one identified neuronal subpopulation (cluster) within that clip. Different clips produced distinct clustering structures and numbers of functional subpopulations, indicating that although neurons recorded during the same video context share broad context-dependent organization, they also exhibit substantial functional diversity within each clip condition. Color scale indicates Spearman correlation values (ρ).

For each cluster, we computed a mean functional map by averaging the neuron–BOLD association patterns of all neurons assigned to that cluster. Most cluster-average maps exhibited strong associations with medial orbitofrontal cortex (mOFC), dorsolateral prefrontal cortex (dlPFC), lateral motor cortex (M1/PM), rhinal cortex, and visual cortical areas V1 and V2. Notably, the direction of these associations varied across clusters, with some exhibiting positive and others negative association patterns (Figure 7). Together, these findings indicate that although shared cinematic context strongly constrains large-scale neuron–BOLD organization, dmPPC neurons nonetheless exhibit substantial functional heterogeneity within the same contextual condition. Rather than forming a uniform population, neurons dynamically organize into multiple distinct functional clusters characterized by different large-scale association patterns across prefrontal, motor, medial temporal, and visual cortical systems.

## DISCUSSION

In this study, we adopted a cross-modality approach to investigate functional diversity within local neuronal populations of the primate dorsomedial posterior parietal cortex (dmPPC). By integrating single-unit electrophysiology with whole-brain fMRI during naturalistic video viewing, we demonstrated that neuron–BOLD functional maps exhibit strong context-dependent organization that differs across cinematic conditions. Compared with neuronal spiking patterns or critical fMRI frames alone, neuron–BOLD functional maps more robustly captured context-dependent structure. Furthermore, although neurons recorded during the same video context exhibited broad similarity in their whole-brain association patterns, neighboring neurons recorded simultaneously from the same electrode still displayed substantial local heterogeneity. Together, our findings support a model in which dmPPC neurons participate in dynamic, context-dependent large-scale brain states rather than fixed functional networks.

Previous studies have examined relationships between electrophysiological activity and BOLD signals, demonstrating that BOLD fluctuations are often more closely associated with local field potentials than with spiking activity alone (Logothetis et al., 2001; Masse et al., 2020; Mukamel et al., 2005; Nir et al., 2008). However, relatively few studies have directly integrated neuronal spiking with whole-brain fMRI to examine how single-neuron activity relates to large-scale cortical organization. Cheng et al. (2023) demonstrated that neuronal and voxel population signals share related information structures through tuning-compatible response correlations, whereas Yang et al. (2026) reported large-scale switching dynamics between inward and outward forebrain activity modes. In contrast to these approaches, our study used neuron–BOLD functional maps as a direct measure for characterizing how individual dmPPC neurons relate to distributed whole-brain activity patterns during naturalistic viewing.

A key finding of the present study is that neuron–BOLD functional maps preserve contextual structure more strongly than neuronal spiking patterns or critical fMRI frames alone. Recent evidence suggests that dmPPC neuronal populations encode temporally evolving contextual information during naturalistic experience and memory retrieval (Zuo et al., 2026), supporting the view that dmPPC contributes to episodic processing through dynamic population-level representations distributed across time and context. In this study, neurons recorded during presentation of the same video sequences exhibited substantially greater similarity in their whole-brain association patterns than neurons recorded during different video contexts. These results suggest that integrating neuronal activity with distributed BOLD dynamics captures aspects of contextual organization that are less apparent when either modality is analyzed independently. Consistent with previous naturalistic-viewing studies, identical video stimuli elicited reproducible large-scale activity patterns across viewers (Hasson et al., 2004, 2008; Kauppi et al., 2010; Kwok & Macaluso, 2015; Meer et al., 2020; Ortiz-Rios et al., 2021; Russ et al., 2021), supporting the interpretation that stimulus-driven cortical organization contributes substantially to the observed neuron–BOLD association structure. In addition, recent work examining cortical organization during natural movie viewing identified parietal functional networks associated with both default mode and action-perception systems (Rajimehr et al., 2024), consistent with the broad context-dependent patterns observed in our dmPPC recordings.

Our findings further suggest that dmPPC neurons participate in distributed cortical systems in a highly context-dependent manner. ROI-based analyses revealed relatively stable association gradients in sensorimotor and cingulate regions, whereas prefrontal and medial temporal regions exhibited substantially greater heterogeneity across video contexts. Such context-dependent organization may reflect the integrative role of dmPPC within distributed higher-order cortical systems (Cavanna & Trimble, 2006; S. Zhang & Li, 2012). fMRI studies have consistently demonstrated strong functional relationships between dmPPC, visual networks, and attention-related regions (Alahmadi, 2021; Hutchinson et al., 2014), while recent work has suggested dimension-specific representations within default mode network structures involved in episodic processing (Procida et al., 2025). Our results extend these observations by suggesting that individual dmPPC neurons exhibit diverse large-scale association patterns that dynamically vary across contextual conditions. In this context, our findings suggest that neuron–BOLD functional maps capture transient, context-dependent association structure during naturalistic viewing rather than stable intrinsic connectivity between neurons and brain regions.

The observed heterogeneity is also consistent with previous evidence that dmPPC neurons exhibit mixed selectivity (Rigotti et al., 2013) for high-dimensional sensory and ethological variables (Wang et al., 2024). Recent electrophysiological studies using ethogram-based behavioral analyses demonstrated that dmPPC neurons encode multiple dimensions of socially relevant information, including foraging, grooming, aggression, and conspecific-related cues (Wang et al., 2024). Our findings complement this work by showing that such neuronal diversity is accompanied by heterogeneous large-scale neuron–BOLD association structure. In addition, higher-order cortical regions are known to exhibit longer temporal receptive windows than early sensory areas (Hasson et al., 2008; Honey et al., 2012), potentially enabling integration of information across extended naturalistic events. The context-dependent association patterns observed in prefrontal and medial temporal regions may therefore reflect dynamic large-scale organization associated with processing temporally extended sensory and cognitive information for episodic memory retrieval (Zuo et al., 2026).

An additional important finding was that neighboring dmPPC neurons exhibited both local similarity and substantial local heterogeneity. Although neighboring neurons were significantly more likely to share similar functional profiles than expected by chance, a substantial subset of simultaneously recorded neurons nonetheless displayed markedly divergent whole-brain association patterns. These findings suggest a nested organizational structure in which shared contextual conditions constrain broad local organization, while neighboring neurons still maintain distinct large-scale association patterns. Similar observations have previously been reported in the inferotemporal cortex using single-unit fMRI mapping approaches (Park et al., 2017; Russ et al., 2023). Our results extend these findings to dmPPC and suggest that functional diversity within local neuronal populations may be a general feature of higher-order association cortex.

Several methodological considerations should be acknowledged. First, the relatively short duration (30 s) of the video clips used to generate clip-specific neuron–BOLD association maps. Although short windows may potentially reduce stability of correlation estimates given the sluggish temporal dynamics of the BOLD signal, our split-half reliability and permutation analyses demonstrated that the clips elicited reproducible stimulus-locked BOLD responses across repeated presentations (Figure S3). Nevertheless, future studies using longer naturalistic sequences may further improve temporal stability and enable investigation of slower context-dependent dynamics. Given the short clip duration, the resulting neuron–BOLD functional maps should be interpreted as transient, stimulus-locked association patterns that emerge during specific naturalistic viewing rather than as stable estimates of intrinsic functional connectivity.

Second, repeated presentation of video clips raises the possibility that increasing familiarity may influence neuronal responses over time. Although repetition is known to modulate neural response magnitude, averaging across repeated trials likely reduced transient trial-specific variability and emphasized stable stimulus-driven components shared across presentations. Third, we cannot exclude the potential influence of unmeasured internal-state variables such as arousal, motivation, attention, or individual differences across animals. Because eye-tracking data were not collected, fluctuations in gaze position or attentional engagement could also contribute to some of the observed variability. Nonetheless, the strong context-dependent similarity observed at the population level suggests that robust stimulus-driven organization emerges despite these uncontrolled factors. Future studies combining multimodal recordings with behavioral and physiological monitoring will be important for further dissociating stimulus-driven and internal-state contributions to neuron–BOLD association structure.

Finally, although the present study emphasizes functional diversity within dmPPC neuronal populations, the relationship between these association patterns and underlying cellular subclasses remains unresolved. Recent studies have demonstrated substantial complexity in the cellular organization of the primate cerebral cortex (Bernard et al., 2012; Chen et al., 2023; Han et al., 2022; Zhang et al., 2025). It is therefore possible that the neuron–BOLD association diversity observed here may partially relate to differences in cell type, microcircuit organization, or long-range connectivity properties. Future work integrating large-scale physiology with cellular-resolution approaches will be necessary to determine how local cellular architecture contributes to large-scale functional organization.

## SUPPLEMENTARY INFORMATION

Supplementary information can be found online.

## ACKNOWLEDGMENTS

The authors thank Makoto Kusunoki for implanting the headposts and the chambers for electrophysiology experiments, and Fabien Balezeau and Marcus Haag for their assistance with awake-monkey fMRI acquisition. We also thank all members of the Kwok lab for their input and support.

## AUTHOR CONTRIBUTIONS

SCK conceived and designed research; XZ, LW, and MO-R. performed research; XZ, LW, SKET, MO-R, and SCK. contributed unpublished reagents/analytic tools; XZ analyzed data. XZ, and SCK drafted and finalized the manuscript with input from LW, SKET, MO-R, DY.

## DATA AVAILABILITY

Original data reported in the study are available in a persistent repository upon publication. DOI/accession number(s): https://osf.io/3cjgb/overview?view_only=50b75691d8cb4dbb899ddc948c28e40b

## Supplementary material

**Figure S1.**
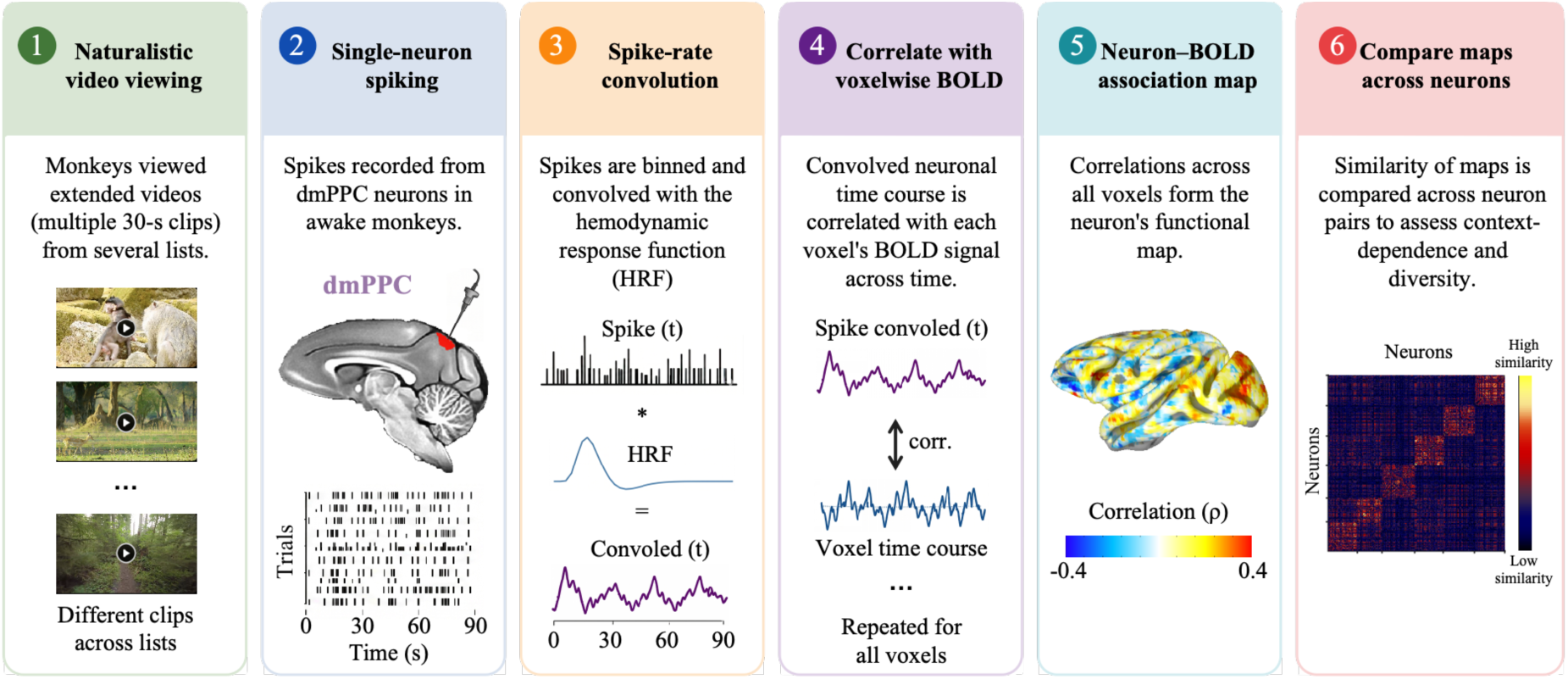
Schematic overview illustrating the full pipeline: From naturalistic video viewing and neuronal recordings to construction and comparison of stimulus-locked neuron–BOLD association maps.

**Figure S2.**
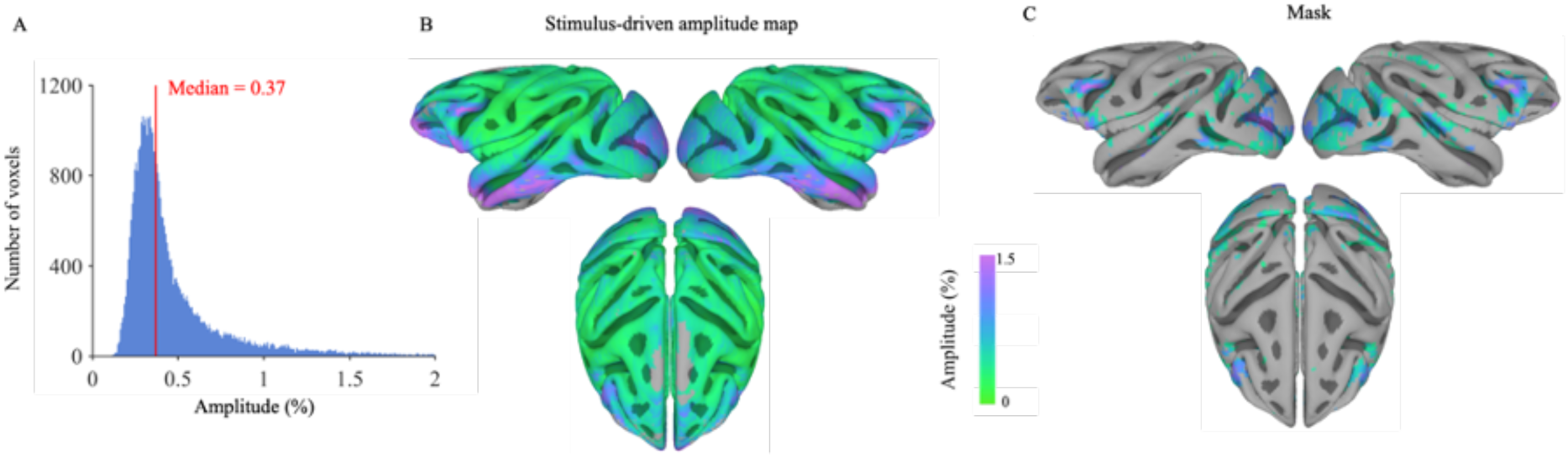
Computation of amplitude-based voxel mask. (A) Distribution of voxel-wise BOLD signal amplitude, quantified as the root mean square (RMS) of the preprocessed BOLD time series and expressed in percent signal change. The red line indicates the median value (0.37%), which was used as the threshold for defining the amplitude mask. (B) Whole-brain map of BOLD signal amplitude (RMS, % signal change), illustrating the spatial distribution of stimulus-driven signal magnitude during video viewing. (C) Amplitude mask obtained by thresholding the RMS amplitude map at 0.37%. Voxels below the threshold were excluded from further analysis. The resulting mask defined the subset of voxels included in the subsequent neuron–BOLD correlation analyses (5027 voxels).

**Figure S3.**
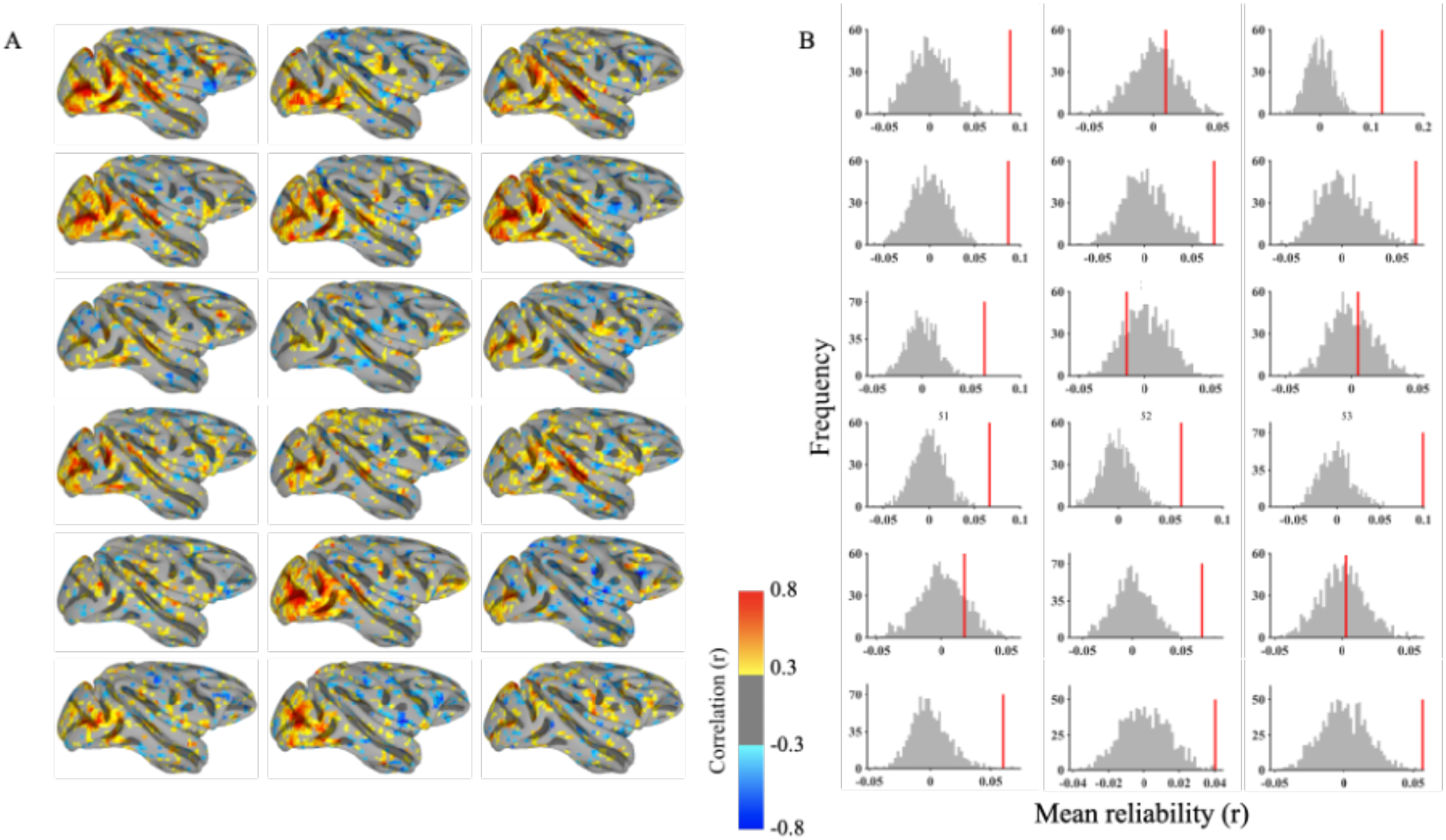
Voxel-wise split-half reliability maps of fMRI signals across 18 video clips. (A) Voxel-wise split-half reliability maps of fMRI signals across 18 video clips. To assess the reproducibility of stimulus-evoked hemodynamic responses within the relatively short (30-s) duration, a split-half reliability analysis was performed for each video clip. Repeated fMRI viewing trials were divided into two independent groups, and the Pearson correlation coefficient (r) was computed between the mean BOLD time courses of these two groups for each voxel. The resulting maps show the degree of temporal consistency across independent presentations. Warmer colors (yellow/red) indicate higher split-half similarity (e.g., r > 0.3), reflecting greater temporal consistency across independent presentations. Reliable responses were particularly prominent in visual and parietal regions, indicating robust stimulus-locked hemodynamic responses during video viewing. (B) Permutation-based statistical evaluation of split-half reproducibility for each video clip. Gray histograms show null distributions of mean reliability generated from 1,000 temporal-shuffling permutations, and red vertical lines indicate the observed reliability values from the original data. For the majority of clips (13 of 18), observed reliability exceeded the 95th percentile of the null distribution, supporting the interpretation that short video clips are sufficient to elicit reproducible, structured BOLD responses.

**Figure S4.**
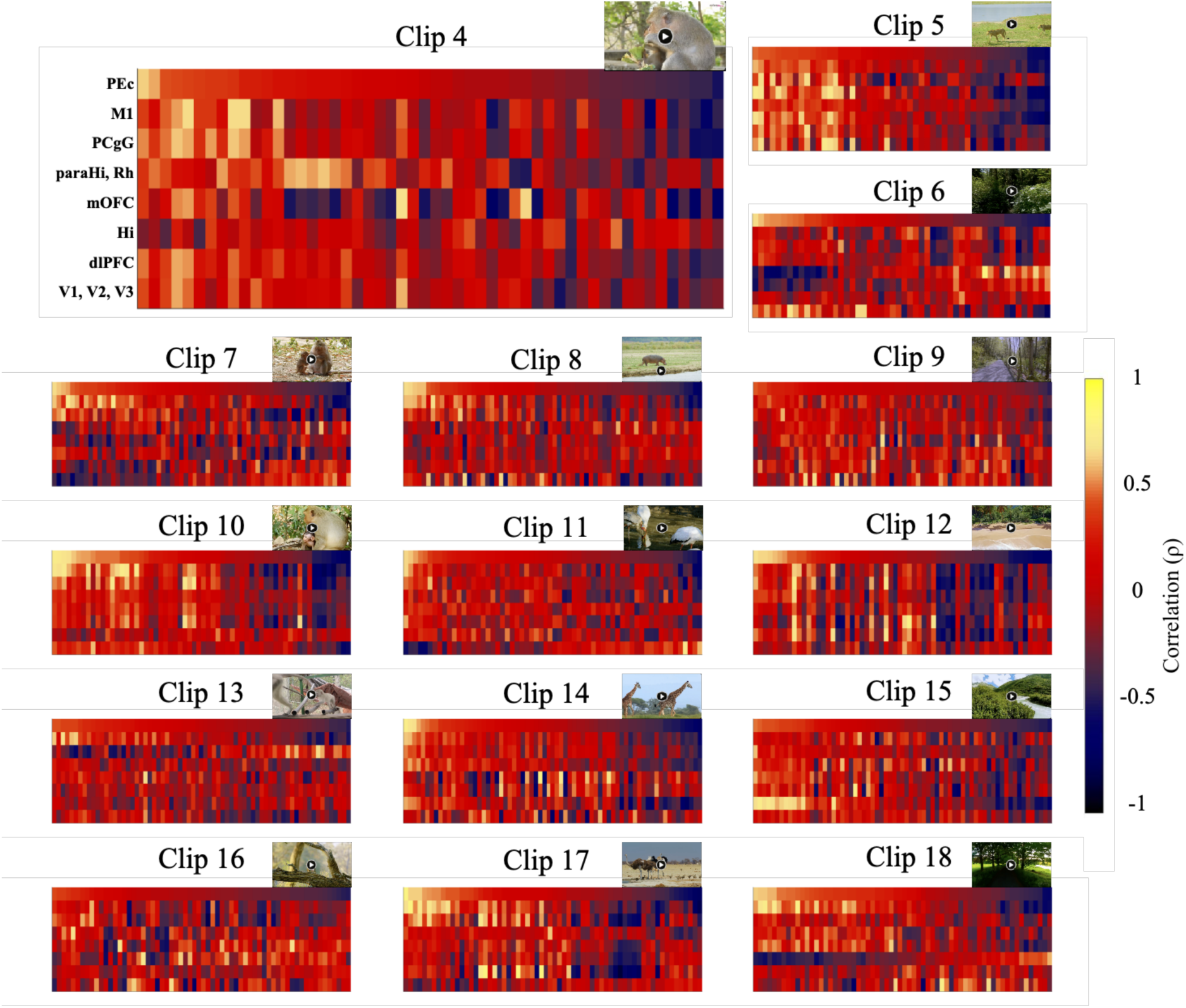
Context-dependent neuron–BOLD association patterns across ROIs during viewing of video clips 4–18. Each heatmap corresponds to one video clip condition (clips 4–18) and shows neuron–BOLD association values across cortical ROIs for all recorded neurons. Columns represent neurons and rows represent ROIs. Neurons are ordered according to their association strength within the dmPPC (PEc; top row), computed as the Spearman correlation between each neuron’s convolved spiking profile and ROI-averaged fMRI BOLD signals. This ordering highlights both shared and divergent association patterns across cortical regions and video contexts. Related to Figure 6.

**Figure S5.**
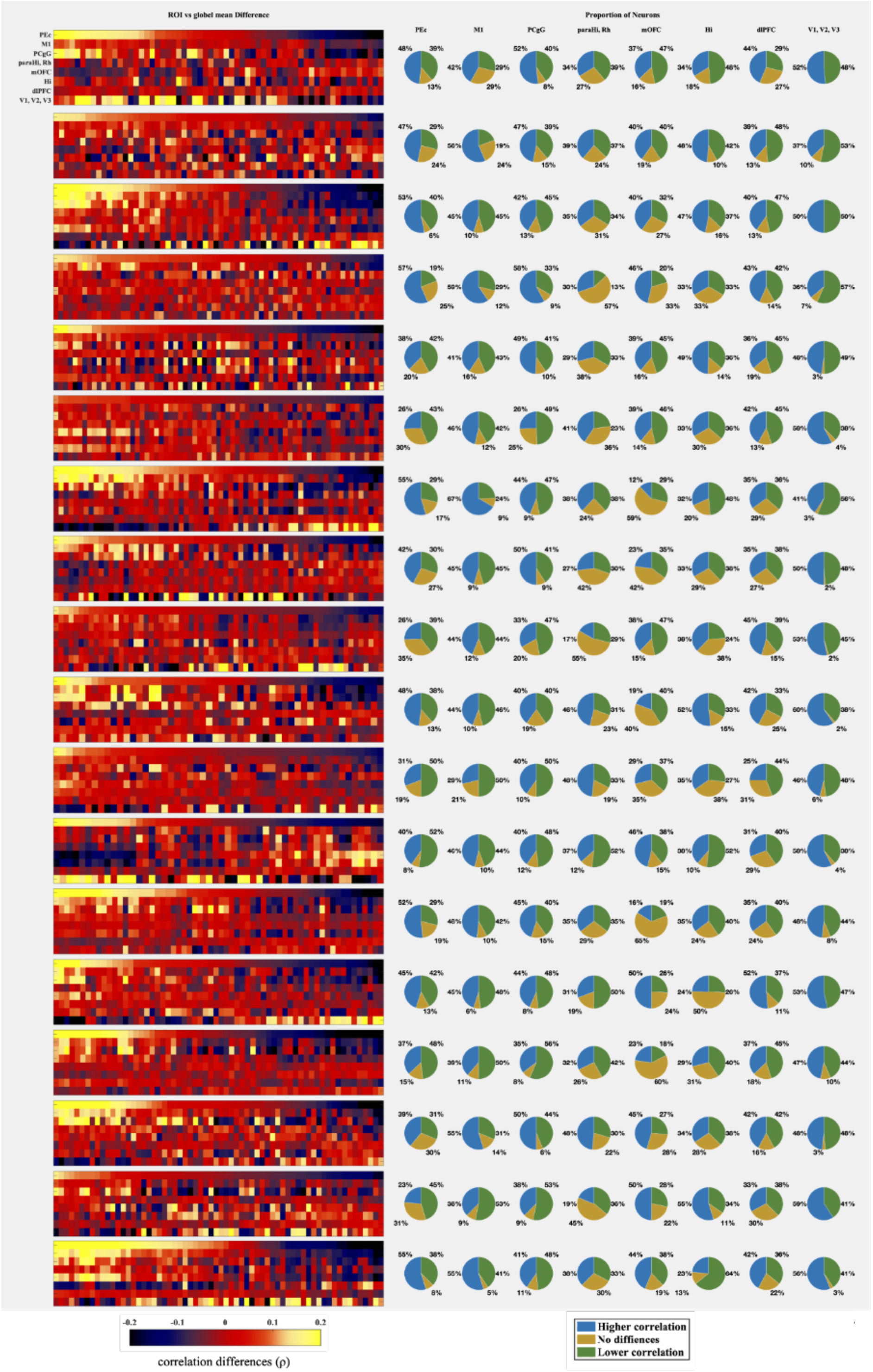
Context-dependent differences between ROI-specific and extra-ROI neuron–BOLD correlations across video clips. Each panel corresponds to one of the 18 video clips. Left: Heatmaps showing, for each neuron, the difference between the neuron–BOLD correlation within a given ROI and the mean neuron–BOLD correlation outside that ROI (extra-ROI correlation). Columns represent neurons and are sorted according to the correlation difference in the dmPPC (PEc; top row). Positive values indicate stronger neuron–BOLD association within the ROI relative to the rest of the brain, whereas negative values indicate relatively weaker association. Right: Pie charts summarizing the proportion of neurons exhibiting significantly higher, lower, or non-significant ROI-specific correlations relative to extra-ROI correlations for each ROI. The analysis shows context-dependent heterogeneity in neuron–BOLD association patterns across cortical regions and video conditions.

**Figure S6.**
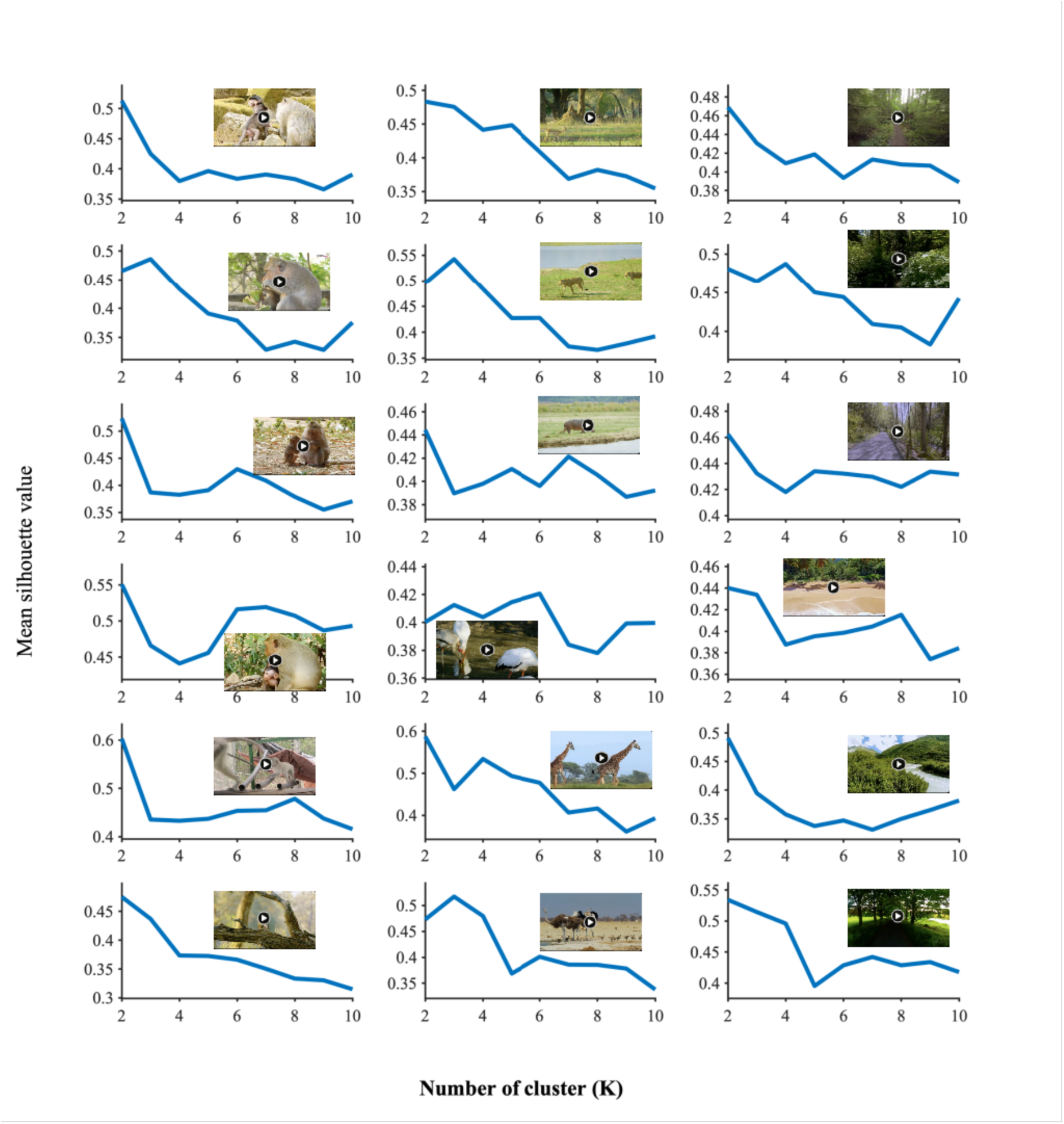
Silhouette analysis of k-means clustering for neuron–BOLD functional maps across video clips. Each subplot corresponds to one of the 18 video clips and shows the mean silhouette value obtained across different numbers of k-means clusters (k = 2–10). Higher silhouette values indicate stronger separation and internal consistency of the resulting functional clusters. The optimal cluster number varied across video contexts, with the highest silhouette scores observed at k = 2 for clips 1, 2, 3, 7, 8, 9, 10, 12, 13, 14, 15, 16, and 18; k = 3 for clips 4, 5, and 17; k = 4 for clip 6; and k = 6 for clip 11. These analyses were used to determine the clustering structure shown in Figure 7.

